# Antimicrobial susceptibility and serotype distribution of *Streptococcus agalactiae* recto-vaginal colonizing isolates from pregnant women at a tertiary hospital in Pretoria, South Africa: an observational descriptive study

**DOI:** 10.1101/564856

**Authors:** Mohamed Said, Yusuf Dangor, Nontombi Mbelle, Shabir A. Madhi, Gaurav Kwatra, Farzana Ismail

## Abstract

**Introduction:** *Streptococcus agalactiae* or Group B Streptococcus (GBS) is a significant cause of neonatal sepsis. Intrapartum antibiotic prophylaxis is recommended for pregnant women identified to be recto-vaginally colonised between 34-37 weeks gestational age to decrease the risk of invasive disease in their newborns. The aim of this study was to investigate serotype distribution and antimicrobial susceptibility patterns of GBS isolates cultured from recto-vaginal specimens during pregnancy.

**Methods:** Sixty-nine archived maternal colonizing isolates were tested against penicillin, erythromycin, clindamycin, vancomycin and levofloxacin. Minimum Inhibitory Concentration (MIC) testing was performed using the E-test method. Serotyping was performed by latex agglutination method.

**Results:** The most common serotypes detected were Ia (54%), III (20%), V (16%), II (6%), IV (2%) and Ib (1%), respectively. All isolates were fully susceptible to penicillin, vancomycin and levofloxacin. Eight (11%) and 50 (56%) isolates showed intermediate resistance to erythromycin and clindamycin respectively, and one isolate was resistant to erythromycin. MLS_B_ phenomenon was noted in 3 (4%) of the isolates.

**Conclusion:** GBS colonizing isolates remain susceptible to penicillin and remains the drug of choice for intrapartum antibiotic prophylaxis and treatment of invasive disease in newbrons. Macrolides should only be used if clinically indicated due to the high prevalence of intermediate resistance. A hexavalent GBS vaccine currently under development would provide coverage for 100% of the isolates identified in this study.

## Introduction

*Streptococcus agalactiae* or Group B Streptococcus (GBS) remains a significant cause of early-onset (<7 days age; EOD) and late-onset (7-89 days age; LOD) invasive disease [1]. The incidence of EOD has declined significantly in countries where universal screening of pregnant women for GBS colonization is undertaken between 34-37 weeks of gestational age and intrapartum antibiotic prophylaxis (IAP) during labour is provided to colonized women [2].

Penicillin remains the drug of choice for IAP and for the treatment of GBS-EOD and LOD. Women with a history of penicillin allergy but at low risk for penicillin anaphylaxis should receive alternative treatment with a cephalosporin such as cefazolin instead of erythromycin or clindamycin [2]. This is due to an increasing resistance of GBS to clindamycin and erythromycin. Reported rates of resistance of GBS to erythromycin range from 25-32% and to clindamycin from 13-20% [2]. Vancomycin is an appropriate alternative for patients with a history of anaphylaxis to penicillin and when an isolate is resistant to clindamycin.

An effective GBS vaccine may prevent a broad scope of GBS associated diseases, such as GBS-EOD, GBS-LOD, spontaneous abortions, stillbirth and maternal bacteraemia [2,4]. One approach of vaccine development is to target the capsular polysaccharide (CPS) of GBS. GBS serological grouping is based on the polysaccharide capsule. There are currently 10 serotypes i.e. Ia, Ib and II-IX. The distribution of the five most common GBS serotypes in South Africa causing invasive disease are III-55.4%, Ia-28.2%, V-7.9%, II-3.6% and Ib-3.4%, II-5% [5]. This compares similarly to the global distribution [6]. Seven to thirty percent of GBS isolates are serologically non-serotypeable [7].

The aim of this study was to determine the serotype distribution of recto-vaginal colonizing isolates from pregnant women and the antimicrobial susceptibility patterns thereof.

## Materials and methods

### Study Design

This was a laboratory based observational study examining 69 archived isolates from a study done in 2014 which investigated the prevalence of GBS colonisation in pregnant women between 26 and 37 weeks gestation [8]. In that study, 284 pregnant women were enrolled from an antenatal clinic and tested for GBS colonisation by Xpert GBS and culture. The colonisation rate was found to be 25% by culture and 24% by Xpert GBS [8]. The GBS isolates were stored in trypticase soy broth with 5% glycerol.

The women had been enrolled and microbiology testing done at the Tshwane Academic Division Microbiology laboratory of the National Health Laboratory Services (NHLS). The serotyping of the isolates was conducted at the Respiratory and Meningeal Pathogens Research Unit (RMPRU, Johannesburg.)

### Specimen processing

The stored isolates were sub-cultured on 5% sheep blood agar and incubated for 24 hours in 5% CO_2_. Beta-haemolytic colonies were then lawned onto Mueller Hinton agar with 5% sheep blood for Minimum Inhibitory Concentration (MIC) testing using Etest (bioMeriuex, France) strips. Five antibiotics were tested for each isolate viz. penicillin, vancomycin, erythromycin, clindamycin and levofloxacin. Plates were incubated for 24 hours in 5% CO_2_ at 35-37°C. The MIC’s were determined using the latest CLSI breakpoints (2015) and the quantitative variables obtained were classified as susceptible, non-susceptible, intermediate and resistant. The MIC’s of GBS isolates which tested non-susceptible or resistant for any of the 5 antibiotics, were repeated and the results confirmed. Furthermore, two observers read the MIC values of all the isolates to minimise inter-observer variability or any form of bias.

As per CLSI guidelines for beta-haemolytic streptococci, MLS_B_ testing was performed on each isolate to test for inducible clindamycin resistance. The isolates were plated on Mueller Hinton plus 5% sheep blood agar, after which erythromycin and clindamycin discs were placed next to each other, 12mm apart. The plates were incubated at 35-37°C in 5% CO_2_ for 18-24 hours. A “D-zone” on the side of the clindamycin disc facing the erythromycin disc was taken as positive for the MLS_B_ resistance phenotype.

Serotyping was performed using the latex agglutination method as described by Kwatra et al [9].

### Ethics Approval

Ethical approval for this study was obtained from the University of Pretoria Faculty of Health Sciences Research Ethics Committee. The ethics reference number is 393/2013.

## Results

The serotype distribution of the 69 isolates were 54% Ia (n=37), 20% III (n=14), 16% V (n=11), 6% II (n=4), 3% IV (n=2) and 1% Ib (n=1).

The antimicrobial susceptibility testing showed 69 (100%) isolates were susceptible to penicillin (MIC range = 0.032-0.125 µg/ml). All isolates were susceptible to vancomycin with 13 (18%) isolates having an MIC at the breakpoint (1μg/ml) (MIC range = 0.38-1µg/ml). Sixty (83%) isolates were sensitive to erythromycin, 8 (11%) isolates were intermediate and 1 (1%) was resistant (MIC range = 0.094-3µg/ml). The erythromycin intermediate isolates belonged to serotypes Ia (3); III (2); IV (1) and V (2).

Thirty (42%) isolates were found to be fully susceptible to clindamycin while 40 (56%) were intermediate-susceptible and no resistant isolates were detected (MIC range = 0.19-0.75 µg/ml). The clindamycin intermediate isolates belonged to serotypes Ia (23), II (1), III (8), IV (1) and V (7). Only 3 (4%; serotypes Ia, III and V) of our isolates displayed a positive MLS_B_ phenotype. All isolates were sensitive to levofloxacin (range = 0.38-1.5 µg/ml).

**Table 1:** Antimicrobial susceptibility of 69 GBS isolates to 5 antimicrobial agents

| Antimicrobial agent | MIC<sub>50</sub> (µg/ml) | MIC<sub>90</sub> (µg/ml) | Range (µg/ml) |
| --- | --- | --- | --- |
| Penicillin | 0.047 | 0.064 | 0.032-0.125 |
| Erythromycin | 0.19 | 0.25 | 0.094-3 |
| Clindamycin | 0.25 | 0.5 | 0.19-0.75 |
| Vancomycin | 0.75 | 0.5 | 0.38 -1 |
| Levofloxacin | 0.5 | 0.75 | 0.38-1.5 |

## Discussion

The current study characterized the antimicrobial resistance patterns in GBS isolates from pregnant women. In this study, sixty-nine (100%) isolates were fully susceptible to penicillin. In a recent Chinese study looking at colonising GBS isolates from pregnant women 100% of isolates were sensitive to penicillin, ceftriaxone, linezolid and vancomycin [10]. Longtin et al. described a case of GBS with reduced susceptibility to penicillin emerging after long term suppressive oral penicillin therapy for a prosthetic joint infection [11].

All isolates in this study were susceptible to vancomycin. There is a paucity of data on vancomycin resistance in GBS isolates, with 2 case reports. These cases involved 2 patients with invasive GBS infection with significant co-morbidities including diabetes, hypertension, congestive cardiac failure, hypercholestrolaemia in 1 patient and end stage renal disease, obesity, cor-pulmonale and chronic osteomyelitis in the 2^nd^ patient [12]. Only one of these patients had previous prolonged exposure to vancomycin. Both isolates were characterised as belonging to serotype II. The vancomycin MIC in both cases were 4ug/ml.

Macrolides are often regarded as alternative therapy for penicillin sensitive patients to treat GBS infections, however resistance to macrolides has increased during recent years in several countries with reported geographical variations [13].In the Japanese study by Matsubara et al (2001) the researchers found much lower rates of resistance to erythromycin and clindamycin, 3% and 1% respectively [14]. In a Malaysian study, 23.3% of isolates were resistant to erythromycin and 17.5% to clindamycin [15].The prevalence of resistance among invasive GBS isolates in the United States ranged from 25%-32% for erythromycin and 13%-20% for clindamycin in reports published during 2006-2009 [2]. Our data suggests a lower level of resistance to these two agents than those observed in the US and is closer to the Japanese data [2,14].

The CLSI recommends MLS_B_ testing for beta-haemolytic streptococci which tests for inducible clindamycin resistance. It was found to be the main mechanism of resistance in GBS isolates isolated from the vagina as well as gastric fluid and ear specimens in a Tunisian study performed by Hrauoui and colleagues [13]. This phenomenon was only noted in 3 (4%) of our isolates. The serotype distribution of these isolates were Ia, III and V.

All isolates in this study were susceptible to levofloxacin. However, there have been reports of fluoroquinolone resistant GBS strains that have emerged in the past decade especially in Asia, including China, Japan and Korea [16]. A study by Wu et al (2017) had confirmed that respiratory samples and elderly patients are two independent risk factors associated with levofloxacin resistance in GBS [16]. In addition the study found that levofloxacin-resistant GBS isolates belonged mainly to the ST19/serotype III serogroup [16].

In low income settings, safe administration of intravenous antibiotics may not always be affordable or feasible, particularly for settings where births do not occur in hospitals. In addition, IAP has not proven to be effective in preventing LOD [1]. Therefore, new strategies for prevention of GBS disease in neonates needs to be considered. Vaccination targeting pregnant women to subsequently protect neonates against GBS infection is a potential option.

Information regarding serotype distribution of GBS strains could guide the development of vaccine candidates. Vaccinating pregnant women against GBS may protect infants from developing invasive GBS disease. Universal screening programs for maternal GBS colonisation followed by IAP in colonised mothers have shown to decrease the incidence of EOD [2]. However, it is thought to have a minimal role in the prevention of LOD. GBS maternal vaccination has the potential to decrease EOD as well impact on LOD.

This study showed that serotypes Ia (54%), III (20%) and V (16%) were the predominant serotypes which is in concordance with other studies conducted among pregnant women in South Africa [5]. Serotypes Ia and III together accounted for 74% of the colonised population in our study, whilst the 3 dominant serotypes accounted for 90% of all cases. These results are in keeping with another South African study which showed that serotype III is the commonest cause of EOD in South Africa, accounting for 41.4% of all cases, whilst serotype Ia accounted for 34.7% of cases [5]. The majority of invasive disease was caused by serotypes Ia, III and V [5]. These 3 serotypes are included in a pentavalent polysaccharide protein conjugate vaccine currently being developed and is in a phase 1 trial [17].

## Conclusions

GBS isolates remain susceptible to penicillin and vancomycin, however, surveillance for resistance needs to be ongoing. Macrolides should only be used once susceptibility results are available as significant rates of intermediate resistance have been detected in these isolates. Ninety percent of colonizing isolates belong to 3 serotypes, viz. Ia, III and V.

## Acknowledgements

I would like to acknowledge Dr Alex Sihlabela from the Department of Obstetrics and Gynaecology at Kalafong Hospital who collected the initial swabs from pregnant women at Kalafong antenatal clinic.

